# Object and object-memory representations across the proximodistal axis of CA1

**DOI:** 10.1101/2020.06.19.160911

**Authors:** Brianna Vandrey, James A. Ainge

**Affiliations:** University of St Andrews, School of Psychology and Neuroscience, St Andrews, Fife KY16 9AZ, United Kingdom

**Keywords:** Place Cell, Hippocampus, CA1 Region, Entorhinal Cortex, Spatial Memory, Episodic Memory

## Abstract

Episodic memory requires information about objects to be integrated into a spatial framework. Place cells in the hippocampus encode spatial representations of objects that could be generated through signalling from the entorhinal cortex. Projections from lateral and medial entorhinal cortex to the hippocampus terminate in distal and proximal CA1, respectively. We recorded place cells in distal and proximal CA1 as rats explored an environment that contained objects. Place cells in distal CA1 demonstrated higher measures of spatial tuning and expressed place fields closer to objects. Further, remapping to object displacement was modulated by place field proximity to objects in distal, but not proximal CA1. Finally, representations of previous object locations were more precise in distal CA1. Our data suggest that lateral entorhinal cortex inputs to the hippocampus support spatial representations that are more precise and responsive to objects in cue-rich environments. This is consistent with functional segregation in the entorhinal-hippocampal circuits underlying object-place memory.

## Introduction

Episodic memory is memory for past personal experiences. Models of the neural circuits underlying episodic memory suggest that spatial input from medial entorhinal cortex (MEC) is combined with non-spatial item information from lateral entorhinal cortex (LEC) to form context-dependent memories within the hippocampus (Ainge, Tamosiunaite, Woergoetter, & Dudchenko, 2007; Ainge, Tamosiunaite, Wörgötter, & Dudchenko, 2012; Ainge, van der Meer, Langston, & Wood, 2007; Eichenbaum, Sauvage, Fortin, Komorowski, & Lipton, 2012; Ferbinteanu & Shapiro, 2003; Hayman & Jeffery, 2008; Leutgeb et al., 2005; Manns & Eichenbaum, 2006). Consistent with this, place cells in the hippocampus encode spatial representations of current and previous object locations (Deshmukh & Knierim, 2013; O’Keefe, 1976) that could be generated by signalling from entorhinal cortex (EC) (Deshmukh & Knierim, 2011; Høydal, Skytøen, Andersson, Moser, & Moser, 2019; Tsao, Moser, & Moser, 2013; Tsao et al., 2018; Wang et al., 2018).

Manipulation studies demonstrate that memory for objects within specific locations is dependent on both EC and hippocampus (Aggleton & Nelson, in press). However, object-location memory can be tested in different ways. In complex tests, multiple objects are presented in different locations and object-location memory is tested by moving and/or replacing them with different objects. Simpler tasks present only two objects and object-location memory is tested by moving or replacing one object to create a new configuration of object and location. Lesions of the hippocampus impair all forms of object-location memory (Barker et al., 2017; Barker & Warburton, 2011; Mumby, Gaskin, Glenn, Schramek, & Lehmann, 2002; Save, Buhot, Foreman, & Thinus-Blanc, 1992; Warburton & Brown, 2010, although see Eacott & Norman, 2004; Langston & Wood, 2010). Manipulations of EC, however, produce a more nuanced deficit. Complete MEC lesions produce deficits in recognising that a familiar object has moved to a novel location (Rodo, Sargolini, & Save, 2017; Van Cauter et al., 2013) and specific inactivation of stellate cells in the superficial MEC has a similar effect (Tennant et al., 2018). In contrast, lesions of LEC impair the ability to remember specific object-location associations (Wilson, Langston, et al., 2013). Object-location memory deficits are more pronounced in both MEC and LEC lesioned animals in more complex tasks that require memory for multiple object-location associations (Kuruvilla & Ainge, 2017; Rodo et al., 2017). These observations demonstrate that the entorhinal-hippocampal network is critical for associating objects with the locations in which they were experienced, and suggests functionally segregated subsystems within the network that integrate object and location information in different ways.

Functional segregation within the entorhinal-hippocampal network is consistent with its anatomy. Inputs from EC are partially segregated in the hippocampus, and projections from LEC and MEC terminate in distinct regions of the CA1 proximodistal axis. LEC sends projections predominantly to distal CA1, bordering the subiculum, while MEC projects predominantly to proximal CA1, bordering CA2 (Masurkar et al., 2017; Naber, Lopes da Silva, & Witter, 2001; Steward, 1976; Tamamaki & Nojyo, 1995; Witter, Wouterlood, & Naber, 2000; Wyss, 1981). Examination of how objects are represented in LEC and MEC also suggests segregated functional networks (Deshmukh & Knierim, 2011, 2013; Høydal et al., 2019; Tsao et al., 2013, 2018). LEC contains cells that develop specific and consistent spatial signals in the presence of objects (Deshmukh & Knierim, 2011, 2013; Tsao et al., 2013, 2018). A subset of these neurons also generate responses to empty positions in which objects have previously been experienced, suggesting a neural correlate for object-location memory (Tsao et al., 2013, 2018). In comparison, a significant proportion of MEC cells encode vector relationships between objects and the position of the animal (Høydal et al. 2019). This is consistent with the suggestion that different types of responses to objects are maintained in functionally separate entorhinal-hippocampal circuits. Further support for this suggestion comes from studies showing distal CA1 is preferentially recruited to process information about objects (Hartzell et al., 2013; Ito & Schuman, 2012; Nakamura, Flasbeck, Maingret, Kitsukawa, & Sauvage, 2013; Nakazawa, Pevzner, Tanaka, & Wiltgen, 2016), and place cells in proximal CA1 demonstrate higher spatial tuning and stability than place cells in distal CA1 in empty environments (Henriksen et al., 2010). However, it is unclear whether differences across the proximodistal axis of CA1 persist in cue-rich environments.

We examined whether place cells in distal and proximal CA1 are differentially modulated by the presence of objects, and whether EC inputs influence the spatial representation of objects in CA1. We recorded place cell activity as rats foraged in an environment that contained objects. We report higher measures of spatial tuning in distal CA1, which receives LEC inputs, in comparison to proximal CA1, which receives MEC inputs. When an object was moved to a new location, remapping responses in distal CA1 were modulated by the proximity of place fields to the displaced object. Further, place fields generated in distal CA1 were more precise for objects and locations where objects were previously experienced. These results suggest that inputs from LEC modulate the precision and object-responsivity of place cells in distal CA1.

## Materials and Methods

### Animals

Animals were adult male Lister-hooded rats (n=7) weighing 330-450g at the time of surgery. Prior to surgery, animals were housed in groups of 2-4 in diurnal light conditions (12-hr light/dark cycle). After surgery, animals were housed individually. All habituation and testing occurred during the light phase. Animals had *ad libitum* access to water throughout the study. To encourage exploration during the behavioral task, animals were food deprived to ≥ 90% of their free-feeding weight. All experiments were conducted under a project license (70/8306) acquired from the UK home office and in accordance with national (Animal [Scientific Procedures] Act, 1986) and international (European Communities Council Directive of 24 November 1986 (86/609/EEC) legislation governing the use of laboratory animals in scientific research.

### Surgical Implantation of Electrodes

Microdrives contained 4 tetrodes, each comprising 4 electrodes. Tetrodes were constructed by twisting together 17 μm platinum-iridium wire. Tetrodes were threaded through a 20-gauge steel cannula, which was secured to the microdrive with dental cement. Each microdrive was fitted with a built-in groundwire and a screw mechanism which could be turned to lower the electrodes vertically into the brain. Before implantation, tetrodes were plated with gold to lower the impedance of the electrode tip to 200-300 kΩ. For surgical implantation of the electrodes, rats were anaesthetised with Isoflurane before being transferred to a stereotaxic frame. The rats were administered an analgesic (Carprofen) subcutaneously prior to incision. The skull was exposed, and the microdrives were implanted aimed at distal (n = 4 animals) or proximal CA1 (n = 2 animals). Where implants were bilateral (n = 1 animal), one microdrive was aimed at each region of CA1. Coordinates for distal CA1 were 5.0 posterior to bregma and 3.2 lateral to midline. Coordinates for proximal CA1 were 3.6mm posterior to bregma and 3.8mm lateral to midline. For each implant, a craniotomy was made at the relevant coordinates, dura was cut, and the electrode was lowered vertically 1.8mm from the surface of the brain. Implants were secured to the skull using a combination of jewellers screws and dental cement. The groundwire of each microdrive was secured to a screw near the front of the skull. Animals were administered oral analgesic (Metacam) in their diet for three days post-surgery.

### Recording

Screening for units commenced within one week after surgery. A recording cable was connected to the microdrive which relayed unfiltered electrical signals from each tetrode to the digital acquisition system. Signals were amplified with a unity-gain operational amplifier, and passed through a pre-amplifier. The signal was bandpass filtered (600-6000 Hz) and amplified (5000-20000 times). To screen for units, the filtered electrical signal for each tetrode was examined for spiking events via an oscilloscope on a computer screen. Further, population-level EEG signal was examined for frequency characteristics of the hippocampus (theta; 8-12 Hz) to infer the position of each electrode in the brain. If no units were detected, electrodes were lowered vertically into the brain at small increments (≥ 50 μm).

### Behavioral Apparatus

The electrophysiology suite included a screening location (a pot lined with a towel) and a test environment. The test environment was a square wooden box (60cm x 60cm x 90cm), with a white floor and black and white vertically striped walls. To secure objects in place within the test environment, square sections of fastening tape were attached to the middle of each quadrant of the box floor. This experiment used an array of junk objects which were approximately the same size as a rat and varied in colour, shape, and texture. Any object used in habituation was not recycled during testing. During behavioral sessions, identical copies of each object were presented across trials. The same copy of each object was used across standard trials. Objects were cleaned thoroughly with veterinary disinfectant before each trial. A local cue (coloured cardboard) was attached to the wall of the upper right quadrant, and stable global cues in the room (eg. lamps) were visible to the animal throughout testing.

### Habituation

Animals were habituated to the electrophysiology suite over five consecutive days prior to surgery. On each day of habituation, each animal was placed in the screening location individually for 10 minutes before exposure to the test environment. On day 1, each animal explored the test environment with their cagemates for 10 minutes. On days 2-5, each animal explored the test environment individually for 10 minutes. On day 5, two identical objects were introduced in the test environment at the locations occupied by objects in the standard trials of the behavioral task. For all trials, the animal was placed in the test environment facing away from the objects.

### Behavioral Task

A behavioral session consisted of five consecutive trials, including two object manipulations bounded by standard trials where the objects were presented in a familiar configuration. In the first trial, the animal encountered two different novel objects in the bottom left and right quadrants of the test environment (Standard Trial 1, S1). In the subsequent trial, the animal encountered a copy of each of the objects from the standard trials, but one was moved to a novel location in the upper left or right quadrant of the environment (Object Displacement, O1). The third trial was a repetition of the standard trial (Standard Trial 2, S2). In the fourth trial, the animal encountered two copies of one object from the standard trial in the bottom right and left quadrants of the test environment. One copy was in a novel configuration of object and location, and one copy was in a familiar configuration (Object-Place Recognition, O2). The final trial was a repetition of the standard trial (Standard Trial 3, S3). Each trial was eight minutes long. The animal rested in the screening location for five minutes between trials. The environment was cleaned with veterinary disinfectant between trials to remove waste and neutralise olfactory cues. Across days, the side on which the object manipulation occurred (left or right) was counterbalanced to be pseudo-random. Video footage of the first three minutes of exploration was recorded by a camera positioned above the environment. After three minutes elapsed, food pellets (Dustless Precision Pellets, 45 mg, BioServ) were scattered randomly throughout the box to encourage exploration of the entire environment.

### Histology

Animals were administered a lethal dose of sodium pentobarbitol and transcardially perfused with phosphate-buffered saline (PBS), followed by 300 ml paraformaldehyde (PFA, 4%). To increase the visibility of the electrode tract, the brain was stored within the skull for 24 hours at 4°C. Brains were then extracted and stored in a 20% sucrose solution prepared in PBS for a minimum of 24 hours at 4°C. The brain was sectioned coronally at 50μm on a freezing microtome. 1:4 sections were mounted on slides and fixed for a minimum of one hour in a paraformaldehyde bath. To counterstain cell bodies, sections were de-fatted with xylene, and rehydrated by briefly immersing the slides in a series of ethanol solutions: 100% ethanol, 50% ethanol solution prepared in distilled water (dH^2^O), then dH^2^O. Slides were then immersed in a cresyl violet solution for two minutes, and washed in running tap water for five minutes. Sections were then dehydrated by briefly immersing the slides in the ethanol solutions in reverse order: dH^2^O, 50% ethanol in dH^2^O, and then 100% ethanol. Sections were then cover-slipped with DPX mountant. To confirm the location of the electrode tracts, slides were examined at a 4x magnification using a light microscope (Leitz Diaplan).

### Behavioral Analysis

Behavioral footage was scored offline. The amount of time spent exploring each object was measured in seconds for all trials. To determine whether the animal preferentially explored the object in a novel spatial configuration, a discrimination ratio was calculated for each object manipulation using the following formula (A. Ennaceur & Delacour, 1988):

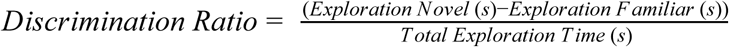

The discrimination ratio is calculated by subtracting the amount of time spent exploring the object in the familiar configuration from the amount of time spent exploring the object in the novel configuration, and then dividing this value by the total exploration time. A positive value indicates an exploratory preference for the object in a novel configuration. For each animal, average discrimination ratios were calculated for each object manipulation. Population means and standard errors of the mean were calculated from these averages.

### Place Cell Identification

Single units were isolated from the raw data using TINT (Axona). First, spike clusters were generated using an automated clustering software, KlustaKwik, which clusters spikes using principal components. Clusters which did not resemble neuronal spikes were removed. The remaining clusters were manually refined by comparing peak amplitude, trough, and time-to-peak and trough on each channel. Only units with a minimum of one place field in any trial of a session, a spatial information score of ≥ 0.5 in all trials where the unit expressed a place field, an average firing rate between 0.1 Hz and 2.5 Hz, and a mean spike duration of ≥ 250 ms were accepted for analysis. To detect place fields for each unit, the position data was sorted into 2 x 2 cm bins. Place fields were defined as contiguous regions of ≥ 6 bins where the firing rate was ≥ 20 % of the peak firing rate for that unit during the trial.

### Quantification of Cluster Quality

To quantify the quality of each cluster, the isolation distance was calculated as described previously (Schmitzer-Torbert, Jackson, Henze, Harris, & Redish, 2005). For each cluster *c* with *n* spikes, isolation distance is defined as the squared Mahalanobis distance of the *n*th closest non-*c* spike to the centre of the cluster. The squared Mahalanobis distance was calculated as:

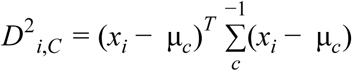

Where *x*_*i*_ is a vector containing feature for spike *i*, and μc is the main feature vector for cluster *c*. High values indicate better isolation. Units with an isolation distance ≥20 were classified as highly isolated, units with an isolation distance ≥10 but <20 were classified as intermediately isolated, and units with an isolation distance <10 were classified as poorly isolated. The calculation of isolation distances required a good connection on all channels of a tetrode. Where a channel was grounded due to noise or disconnection, cluster quality was manually categorised as high, intermediate, or poor by visual comparison against clusters for which an isolation distance value could be determined. Where the same unit was recorded across multiple consecutive days, the recording with the highest average spatial information score was included in the analysis, and other recordings of this unit were discarded. Repeat recordings were determined by examining the shape of the waveform, the tetrodes on which it was recorded, and location of the place field(s).

### Analysis of Place Cell Characteristics

Isolated units were processed offline using customised MATLAB scripts. Rate maps were generated by dividing the area of the box into pixels corresponding to 2 x 2 cm bins of the environment. The firing rate in each pixel was determined by dividing the number of spikes by the dwell-time of the animal in that bin. Firing rate maps depict the firing rate of each bin in colour, where blue represents the lowest firing rate and red represents the highest firing rate. The firing rate maps were analysed to extract the following characteristics: spatial information content, selectivity, spatial coherence, average firing rate, peak firing rate, place field frequency, and place field size.

The spatial information content of a unit, presented as a ratio of bits/spike, indicates the amount of information about the location of an animal which is encoded in each spike. This was calculated using the following formula (Skaggs, McNaughton, & Gothard, 1993):

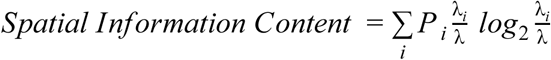

Where λ_i_ is the average firing rate of a unit in the *i-*th bin, λ is the overall average firing rate, and *p*_i_ is the probability of the animal being in the *i*-th bin (dwell time in the *i*-th bin / total recording time). The average firing rate was calculated by dividing the total number of spikes in a trial by the trial duration, and the peak firing rate was the maximum firing rate within the firing field(s) of the cell. Selectivity is a measure of how specific the spikes from the cell are to the place field(s) in an environment and was calculated by dividing the maximum firing rate by the average firing rate. Spatial coherence estimates how coherent a firing field is by determining the extent to which firing rates within a pixel are matched with firing rates in adjacent pixels. This measure is calculated by correlating firing rates within each pixel with the firing rates in eight adjacent pixels, and returning the z-transform of this correlation (Muller & Kubie, 1987). To determine the stability of rate maps across trials, each pixel of the rate map from one trial was correlated with the corresponding pixel from a rate map from a second trial, generating a Pearson’s correlation coefficient. Pixels corresponding to locations in the environment which the animal did not visit in either trial were discarded.

### Analysis of Place Fields

To examine the location of place fields across the environment, the area of the test environment was divided into 8 × 8 bins and the coordinates assigned to the centroid of each place field were plotted across these bins. The centroid of a place field was defined as the average position of the pixels of a place field along the X and Y axis, weighted by the firing rate of those pixels. Using this division of the environment, each quadrant of the environment constituted an array of 16 bins, where the four inner bins (15 x 15 cm) correspond to the location of the object in each quadrant and the outer 12 bins correspond to locations around the object within the quadrant. Frequencies of place fields in each quadrant, and at previous and current object locations were extracted using these criteria. To determine the distance of place fields from objects in the environment, the Euclidean distance between the object centroid, defined as the centre of the object quadrant, and the place field centroid, was measured.

### Analysis of Remapping and Trace Firing

Population changes in spatial coding were quantified by examining the correlation of rate maps across trials. Remapping of individual cells in response to object displacement was quantified by examining the location of place field centroids and correlation of rate maps across S1, O1, and S2. Any cell which expressed no field in S1 and O1 or had a correlation coefficient between S1 and O1 which was greater than or equal to the average correlation coefficient between S1 and S2 for that region (distal or proximal) was categorised as non-remapping. The place fields of the remaining cells were examined for patterns corresponding to remapping behaviors that have been described previously (Deshmukh & Knierim, 2013; Lenck-Santini, Rivard, Muller, & Poucet, 2005; Manns & Eichenbaum, 2009; Muller & Kubie, 1987). Place cells were categorised as remapping if they expressed a place field in the novel object quadrant in O1, but not S1, or the peak firing rate within a pre-existing place field in the novel object quadrant was reduced ≥ 25% in O1. Place cells were also categorised as remapping if a new place field appeared at any location in the environment, if the number of place fields reduced between S1 and O1, or if a pre-existing place field shifted ≥ 7.5 cm. 7.5 cm was chosen as cut-off value given that this distance corresponds with the widths of the bins used to generate the plots of centroid locations. Remapping of place field locations was not examined for the novel object-place recognition trial given that the literature does not predict remapping to this type of manipulation (Deshmukh & Knierim, 2013; Lenck-Santini et al., 2005; Manns & Eichenbaum, 2009).

To quantify rate remapping in response to a change in object identity, firing rate changes were calculated as the normalised rate differences between the first standard trial and the object-place recognition trial (O2) and the second standard trial and O2. These values were calculated by taking the absolute value of the difference in firing rate between the two trials, divided by the sum of the firing rates across the two trials (Lu et al., 2013).

Trace firing was quantified by examining the location of place field centroids across S1, O1, and S2. Place cells were categorised as ‘misplace’ cells if they expressed a place field in the empty quadrant which previously contained the displaced object in O1, but not S1 (O’Keefe, 1976). Place cells were categorised as remap ‘trace’ cells if they expressed a place field in the quadrant which contained the displaced object in O1, but not S1, and a place field persisted in this quadrant in S2. Place cells were categorised as non-remap ‘trace’ cells if they did not express a place field in the quadrant which contained the displaced object in S1 or O1 but did express a place field in this quadrant in S2.

### Statistical Analyses

All statistics were calculated in SPSS (IBM, version 24). To determine whether there was a significant difference across groups for behavior, place cell characteristics, and vector distances, univariate ANOVAs were conducted with electrode location (distal versus proximal) as a between-subjects factor. For place cell characteristics, this analysis was conducted using average values collapsed across standard sessions alone and object manipulation sessions alone. To determine whether patterns of remapping or trace firing in response to changes in the position of objects were observed at similar proportions across groups, observed frequencies were compared across proximal and distal CA1 using a Chi-Square test of independence.

## Results

We recorded from place cells in distal and proximal CA1 as rats foraged in a square environment containing two different objects (Figure 1A). Object locations were consistent in standard trials (S1, S2, S3). We also examined place cell responses to changes in object location (O1) or object identity (O2). The time spent exploring the object in a novel configuration was measured in each manipulation trial to confirm recognition of the spatial change (Figure S1).

**Figure 1:**
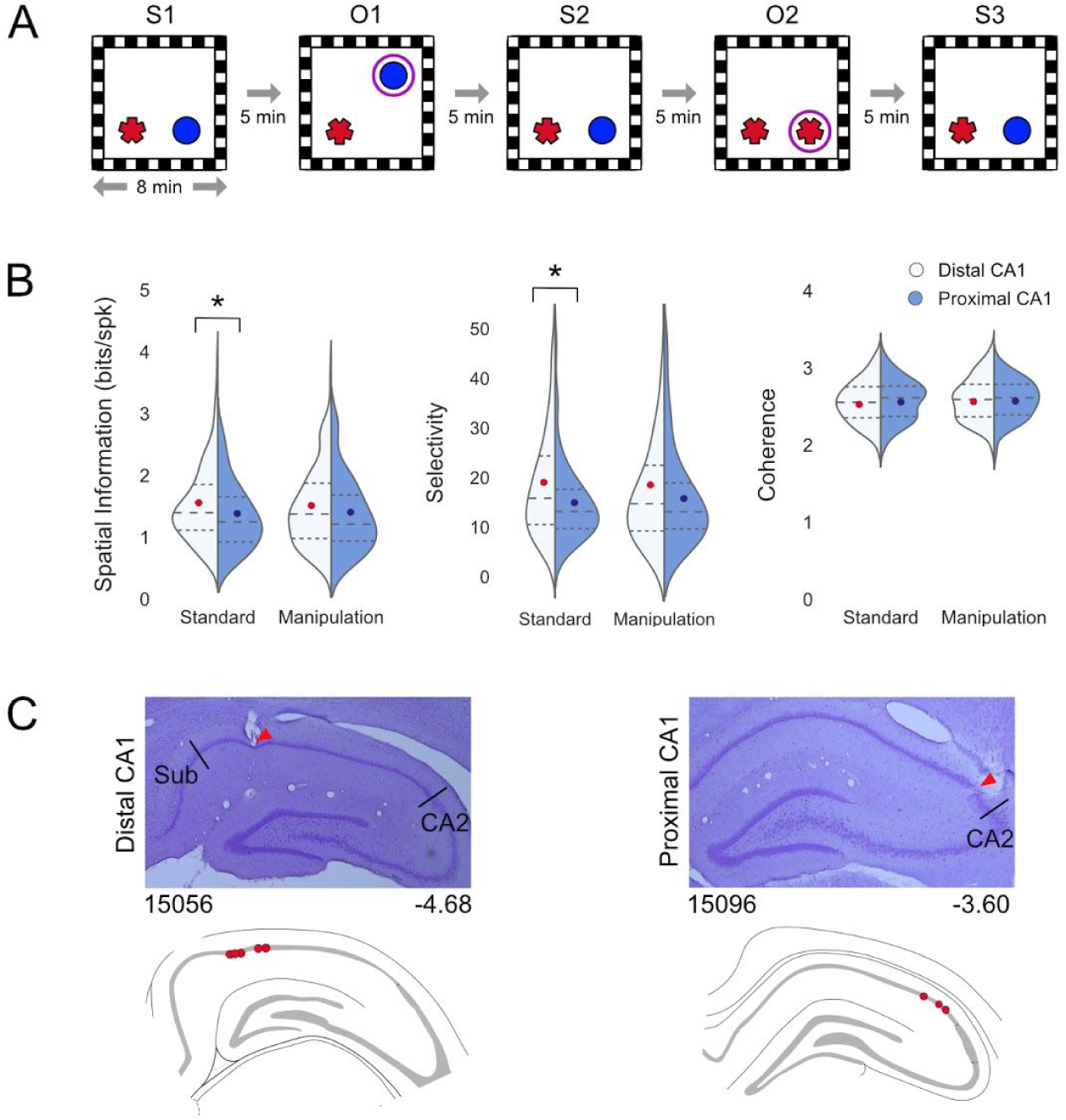
Spatial tuning across the proximodistal axis of CA1. A) Schematic of test trials. Each test session consisted of three standard trials (S1, S2, S3) interleaved with two manipulation trials where an object was displaced to a novel location (O1) or appeared in a novel object-place configuration (O2). Circles indicate the novel configuration in the manipulation trials. B) Violin plots comparing spatial information (left), selectivity (middle), and spatial coherence (right) across place cells in distal (light grey) and proximal CA1 (violet). Width of plot indicates relative frequency of values. Dotted lines indicate upper interquartile range, median, and lower interquartile range. Coloured dots represent mean values (distal = red, proximal = blue). Asterisk indicates *p-*value < 0.05. C) Representative examples of electrode tracts in distal (left) and proximal CA1 (right) in coronal sections of tissue stained with cresyl violet. Red arrow indicates tract location. Numbers indicate animal (left) and caudal distance from bregma (right). Schematic beneath each image shows electrode tract locations for all animals in each group. Each red dot indicates a single animal.

We first asked whether the increased spatial tuning in proximal CA1 relative to distal CA1 reported in empty environments (Henriksen et al., 2010) persists in an environment that contains objects. We examined spatial tuning in distal and proximal CA1 in standard and manipulation trials (1606 place fields [distal n = 1305, proximal n = 301] from 292 units [distal n = 238, 5 animals; proximal n = 54, 3 animals]; Figure 1B-C). Cluster qualities were similar across distal and proximal CA1 (highly isolated, distal: 111/238 cells, 46.6%, proximal: 22/54 cells, 40.7%, *χ*^*2*^ (1) = 0.617, *p* = 0.432; intermediately isolated: distal: 102/238 cells, 42.9%, proximal: 25/54 cells, 46.3%, *χ*^*2*^ (1) = 0.2118, *p* = 0.645, poorly isolated: distal: 25/138 cells, 10.5%, proximal: 7/54 cells, 13.0%, *χ*^*2*^ (1) = 0.273, *p* = 0.602). Distal CA1 place fields contained more spatial information than proximal CA1 place fields in standard trials (*F*_(1, 290)_ = 3.941, *p* = 0.048, η^2^ = 0.013), but not manipulation trials (*F*_(1, 290)_ = 1.465, *p* = 0.227). Further, place cell selectivity in distal CA1 was higher in standard trials (*F*_(1, 290)_ = 5.830, *p* = 0.016, η^2^ = 0.020), but not manipulation trials (*F*_(1, 290)_ = 1.478, *p* = 0.225). Spatial coherence in both regions was similar in standard (*F*_(1, 290)_ = 0.208, *p* = 0.648) and manipulation trials (*F*_(1, 290)_ = 0.002, *p* = 0.966). These findings contrast previous reports of higher spatial tuning in proximal CA1, which receives inputs from MEC, in environments devoid of local cues (Henriksen et al., 2010). Our data demonstrate that prominent and stable objects drive increased spatial tuning in distal CA1 place cells that receive inputs from LEC.

Place cells in distal CA1 express more place fields than place cells in proximal CA1 in an empty environment (Henriksen et al., 2010). However, objects modulate the size and frequency of place fields in distal CA1 (Burke et al., 2011), indicating that a different pattern of place field expression could emerge in our environment. We observed no difference in the number or size of place fields across the proximodistal axis (Figure 2A). The number of place fields expressed was similar in both regions of CA1 in standard (*F*_(1, 836)_ = 1.353, *p* = 0.245) and manipulation trials (*F*_(1, 563)_ = 0.081, *p* = 0.777), and although the place fields expressed by proximal CA1 place cells were smaller in all trials, this was not significant. Our data show no gradient in the size or frequency of place field expression across the proximodistal axis of CA1 in environments that contain objects.

**Figure 2:**
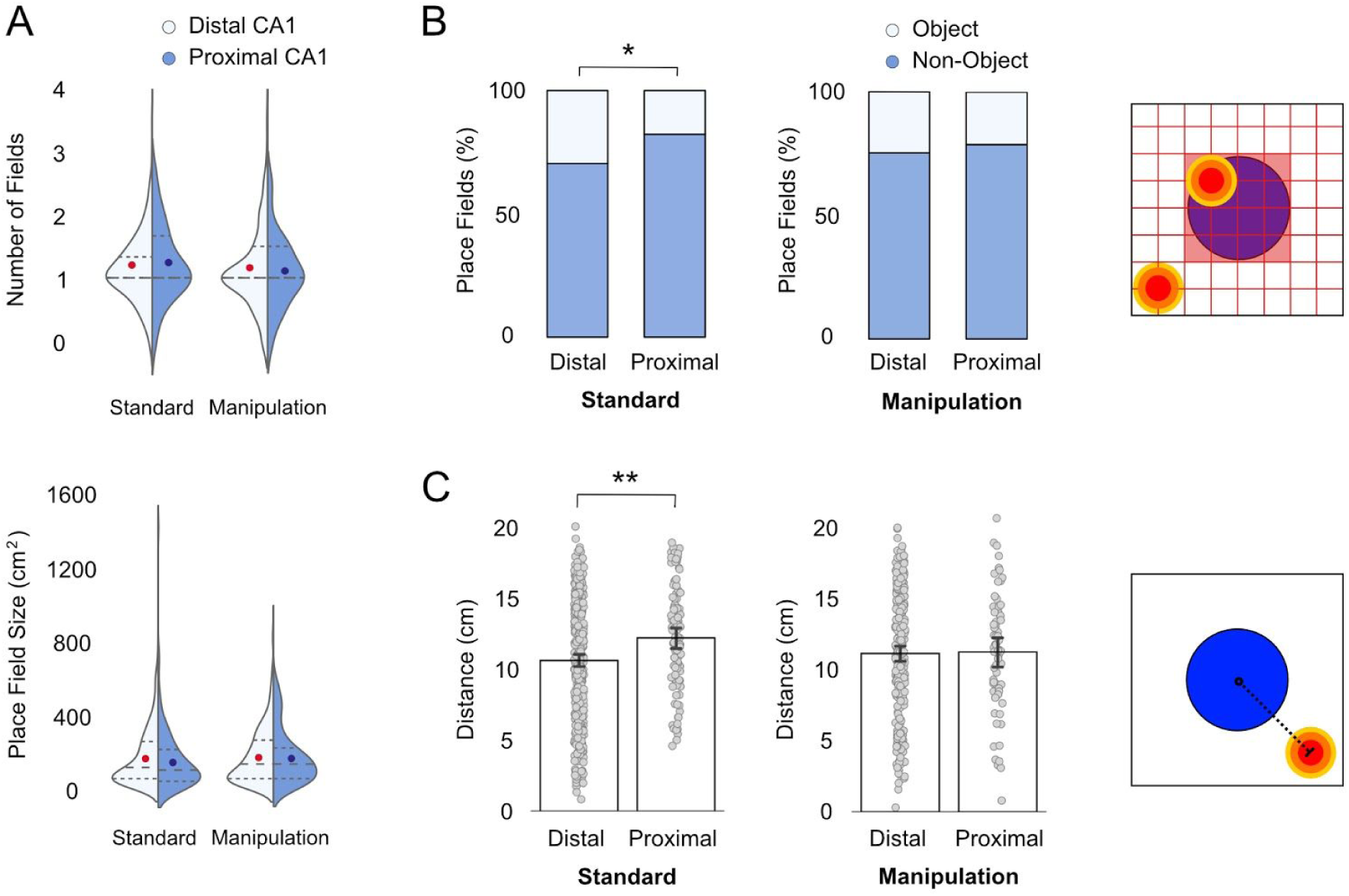
Place field frequency and location across the proximodistal axis of CA1. A) Violin plots comparing place field frequency (top) and size (bottom) across place fields expressed by place cells in distal (left, light grey) and proximal CA1 (right, violet). Width of plot indicates relative frequency of values. Dotted lines indicate upper interquartile range, median, and lower interquartile range. Coloured dots represent mean values (distal = red, proximal = blue). B) (Left) Stacked bar plots comparing the proportion of place fields that are expressed in bins of the environment that correspond to the object (light grey) or at other locations in the quadrant (violet) in standard (left) and manipulation trials (right). Data is shown only from fields expressed in quadrants of the environment which contain an object. (Right) Schematic shows a quadrant of the environment that contains an object with representative place fields that would be categorised as ‘object’ and ‘non-object’. The grid indicates the division of the quadrant into 16 bins, each of which are 7.5 × 7.5 cm. The middle 4 bins correspond to the object position. C) (Left) Bar graphs compare the distance of place field centroids from the center of the object in standard (left) and manipulation trials (right). Each grey dot represents a single field. Asterisk indicates *p-*value < 0.05 (*) or < 0.01 (**). (Right) Schematic shows a quadrant of the environment that contains an object and a place field. The distance of the place field centroid from the object is calculated as the euclidean distance between the centroid and the centre of the quadrant. This distance is represented by a dotted black line.

Given our observation of higher spatial tuning in distal CA1, we next asked whether distal CA1 place fields represent object locations more accurately than proximal CA1 place fields. The positions of place field centroids were analysed in relation to object locations for all place fields expressed within quadrants of the environment that contained an object. The centroid positions of distal CA1 place fields were more likely to correspond to the object location in standard trials (Figure 2B; *χ*^*2*^ (1) = 5.673, *p* = 0.017), but not manipulation trials (*χ*^*2*^ (1) = 5.673, *p* = 0.462) relative to proximal CA1 place fields. In addition, distal CA1 place fields were nearer to the objects than proximal CA1 place fields in standard trials (Figure 2C; *F*_(1, 471)_ = 12.044, *p* = 0.001, η^2^ = 0.025), but not manipulation trials (*F*_(1, 307)_ = 0.021, *p* = 0.885). This demonstrates that place cell representations of quadrants that contain objects are more spatially tuned to object locations if they receive inputs from LEC rather than MEC. However, quantification of the proportions of cells that encode object locations revealed that place cells in proximal CA1 expressed a higher proportion of their place fields in the object quadrants in standard trials (*χ*^*2*^ (1) = 3.862, *p* = 0.049) and manipulation trials (*χ*^*2*^ (1) = 4.07, *p* = 0.044). This may represent an influence of signalling from object-vector cells in MEC (Høydal et al., 2019). Our data demonstrate differences in object representations within entorhinal-hippocampal networks. Proximal CA1 place cells are more likely to fire in the vicinity of objects, yet distal CA1 place cells more precisely encode object locations when object positions are stable.

We next asked whether differential input from EC modulates place field stability in CA1. Our observation of increased spatial tuning in distal CA1 was matched by higher stability of place cells in distal CA1 across trials (Figure 3A-B). We compared correlations between firing rate maps from the first standard trial and all subsequent standard trials and although the correlations in distal CA1 were systematically higher than in proximal CA1, this difference only reached significance for the comparison with object displacement trial (*F*_(1, 290)_ = 8.420, *p* = 0.004, η^2^ = 0.028). These observations indicate that place cells in distal CA1 maintain more stable spatial representations of environments that contain objects, particularly when the location of an object changes.

**Figure 3:**
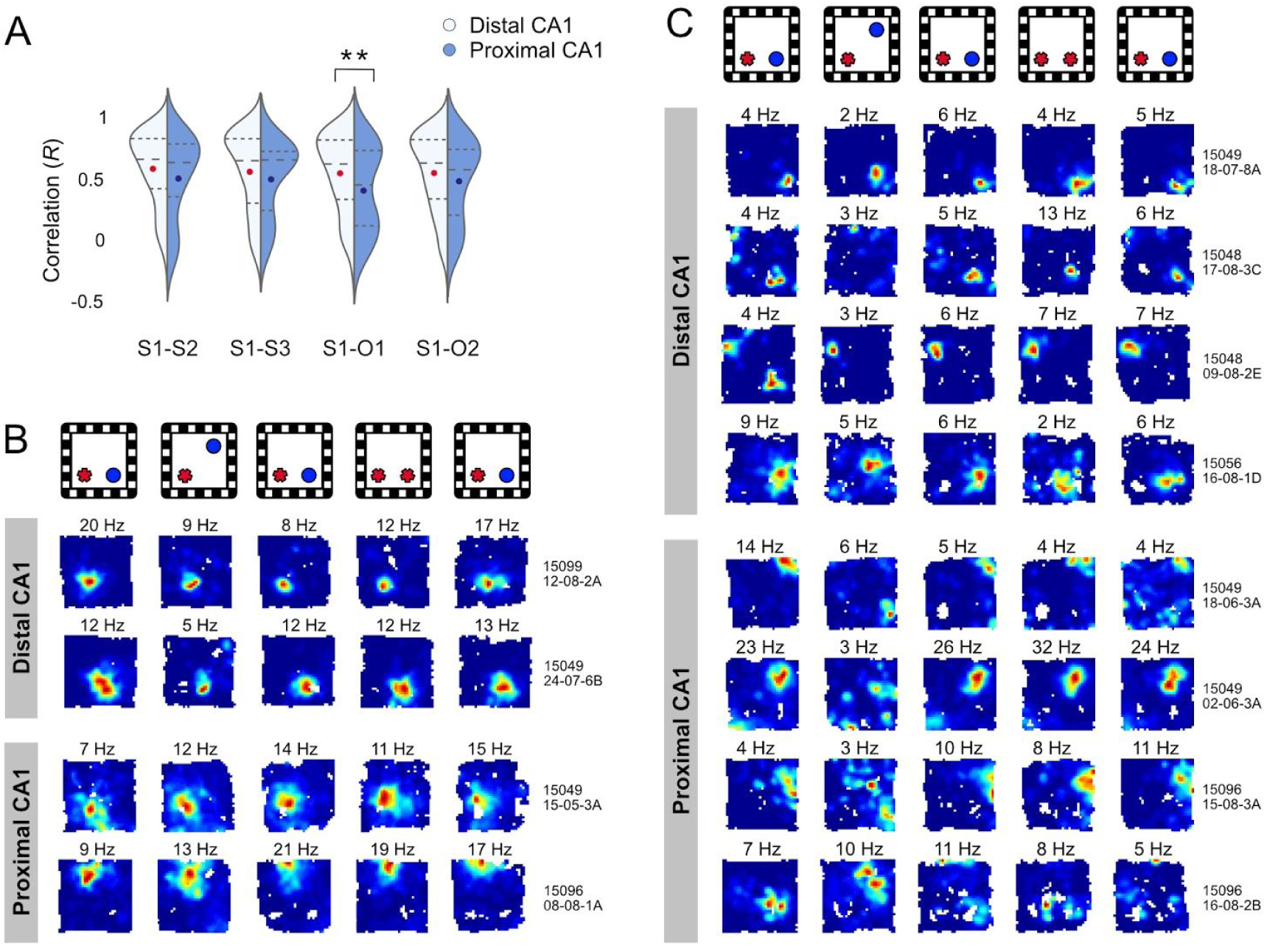
Remapping across the proximodistal axis of CA1. A) Violin plots comparing the stability of place cells across the first and second standard trial (S1-S2), first and last standard trial (S1-S3), first standard trial and object displacement (S1-O1), and first standard trial and object-place (S1-O2). Stability is quantified as the correlation between firing rate maps across trials for each cell, as calculated using Pearson’s product-moment coefficient *(R*). Each plot shows distribution of values for place cells in distal CA1 (left, light grey) and proximal CA1 (right, violet). Width of plot indicates relative frequency of values. Dotted lines indicate upper interquartile range, median, and lower interquartile range. Coloured dots represent mean values (distal = red, proximal = blue). Asterisks indicate *p*-value < 0.01 (**). B) Examples of cells that do not remap in distal (top) and proximal CA1 (bottom). Representative examples are shown that have a correlation value within one standard deviation of the population mean for that region across S1 and O1. Warm colours indicate high firing rates, and cool colours indicate low firing rates or no firing. Peak firing rates for each trial are indicated above the rate maps. C) Examples of remapping place cells in distal (top) and proximal CA1 (bottom). Representative examples are shown that have a correlation value within one standard deviation of the population mean for remapping cells in that region across S1 and O1.

Place cells in the hippocampus remap when objects are moved in an environment by changing the expression of their place fields (Deshmukh & Knierim, 2013; Lenck-Santini et al., 2005; Manns & Eichenbaum, 2009; Muller & Kubie, 1987), but it is unclear how these responses are generated. We therefore asked whether remapping responses to object displacement in CA1 are driven by EC inputs. The proportion of place cells that remapped in the object displacement trial was similar in distal and proximal CA1 (*χ*^*2*^ (1) = 1.326, *p* = 0.250), and the patterns of remapping conformed to those reported previously (Deshmukh & Knierim, 2013; Lenck-Santini et al., 2005; Muller & Kubie, 1987) (Figure 3C). Previous studies have shown that a place cell is more likely to remap when an object is displaced if it expresses a place field near the object before it is moved (Lenck-Santini et al., 2005). Our observation of more precise representations of object positions in distal CA1 suggests that remapping in this region could be driven by proximity of place fields to objects in the standard trials. To examine this possibility, we categorised place cells that expressed place fields in the first standard trial as ‘near’ the object if a place field centroid was located in the quadrant containing the object which would undergo the manipulation, or ‘far’ from the object if all place field centroids were located in the other quadrants. The proportion of cells that remapped in response to object displacement was higher in ‘near’ place cells in distal CA1 (Figure 4A; *χ*^*2*^ (1) = 7.0856, *p* = 0.008), but not proximal CA1 (*χ*^*2*^ (1) = 1.182, *p* = 0.277). Consistent with this finding, the correlations between rate maps across the first standard trial and object displacement were significantly lower for ‘near’ place cells in distal CA1 (Figure 4B; *F*_(1, 196)_ = 10.225, *p* = 0.002, η^2^ = 0.050), but not proximal CA1 (*F*_(1, 42)_ = 0.282, *p* = 0.598). These data demonstrate that place cells in both distal and proximal CA1 encode changes in object position. However, the remapping responses to object displacement are modulated by the proximity of place fields to the object in cells that receive inputs from LEC.

**Figure 4.**
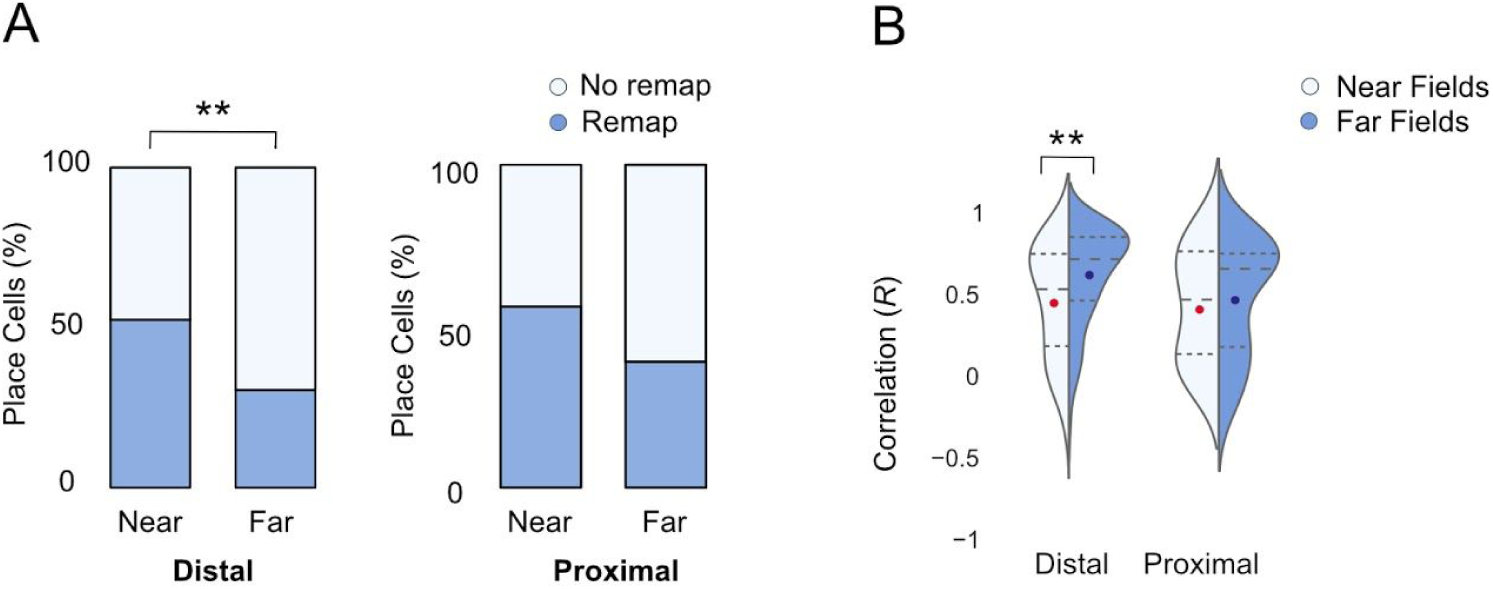
Remapping in near and far fields across the proximodistal axis of CA1. A) Stacked bar plots showing the proportion of remapping cells in distal and proximal CA1 that expressed place fields in the first standard trial. Cells were categorised as ‘near’ if they expressed a place field in the object quadrant before it was displaced, and ‘far’ if they only expressed place fields elsewhere in the environment. B) Violin plot that compares the correlation between rate maps across the first standard trial and object displacement for place cells with fields near the displaced object (light grey) and away from the displaced object (violet) for distal (left) and proximal CA1 (right). Asterisks indicate *p*-value < 0.01 (**).

Previous studies report that changes in object identity are reflected by changes in firing rate rather than the remapping of place fields (Komorowski & Manns, 2009; Larkin, Lykken, Tye, Wickelgren, & Frank, 2014; Manns & Eichenbaum, 2009). LEC neurons encode conjunctive information about object positions in their firing rate (Keene et al., 2016) and LEC lesions alter rate remapping to changes in context in the hippocampus (Lu et al., 2013). These observations raise the possibility that rate coding of novel object-place configurations is stronger in distal CA1. However, firing rates did not vary as a function of trial in distal or proximal CA1, and differences in firing rate between the first or second standard trial and object-place recognition trial were similar across regions (Figure S2). These data demonstrate that place cells do not show robust rate remapping to changes in object identity in CA1.

A striking feature of the hippocampus is that a subset of place cells fire at empty locations where an object was previously located (Deshmukh & Knierim, 2013; O’Keefe, 1976). These cells bear similarity to LEC ‘trace’ cells (Tsao et al., 2013, 2018) and might represent a neural mechanism for object-place memory. We examined trace firing across the proximodistal axis of CA1 to determine whether the characteristics of trace cells differ depending on inputs from EC. Trace cells were observed in similar proportions across distal and proximal CA1 (Figure 5A-B; *χ*^*2*^ (1) = 0.396, *p* = 0.529). A subset of trace cells expressed place fields in the newly empty quadrant in the object displacement trial, consistent with the ‘misplace’ cells described by O’Keefe (O’Keefe, 1976). Misplace cells were observed at similar frequencies across distal and proximal CA1 (*χ*^*2*^ (1) = 0.503, *p* = 0.478). A second subset of trace cells expressed a place field in the novel quadrant, consistent with the ‘object-place memory’ cells described by Deshmukh & Knierim (2013). Object-place memory cells were observed at similar frequencies across distal and proximal CA1 (*χ*^*2*^ (1) = 1.018, *p* = 0.313), and could be further divided into two groups. Some object-place memory cells immediately remapped to express a place field in the novel quadrant when the object was moved, and persisted firing in that quadrant in subsequent trials after the object was returned to its original location (‘remap trace’ cells). However, some cells expressed a new place field in the empty novel quadrant, but only after the object was returned to its original location (‘non-remap trace’ cells).

**Figure 5:**
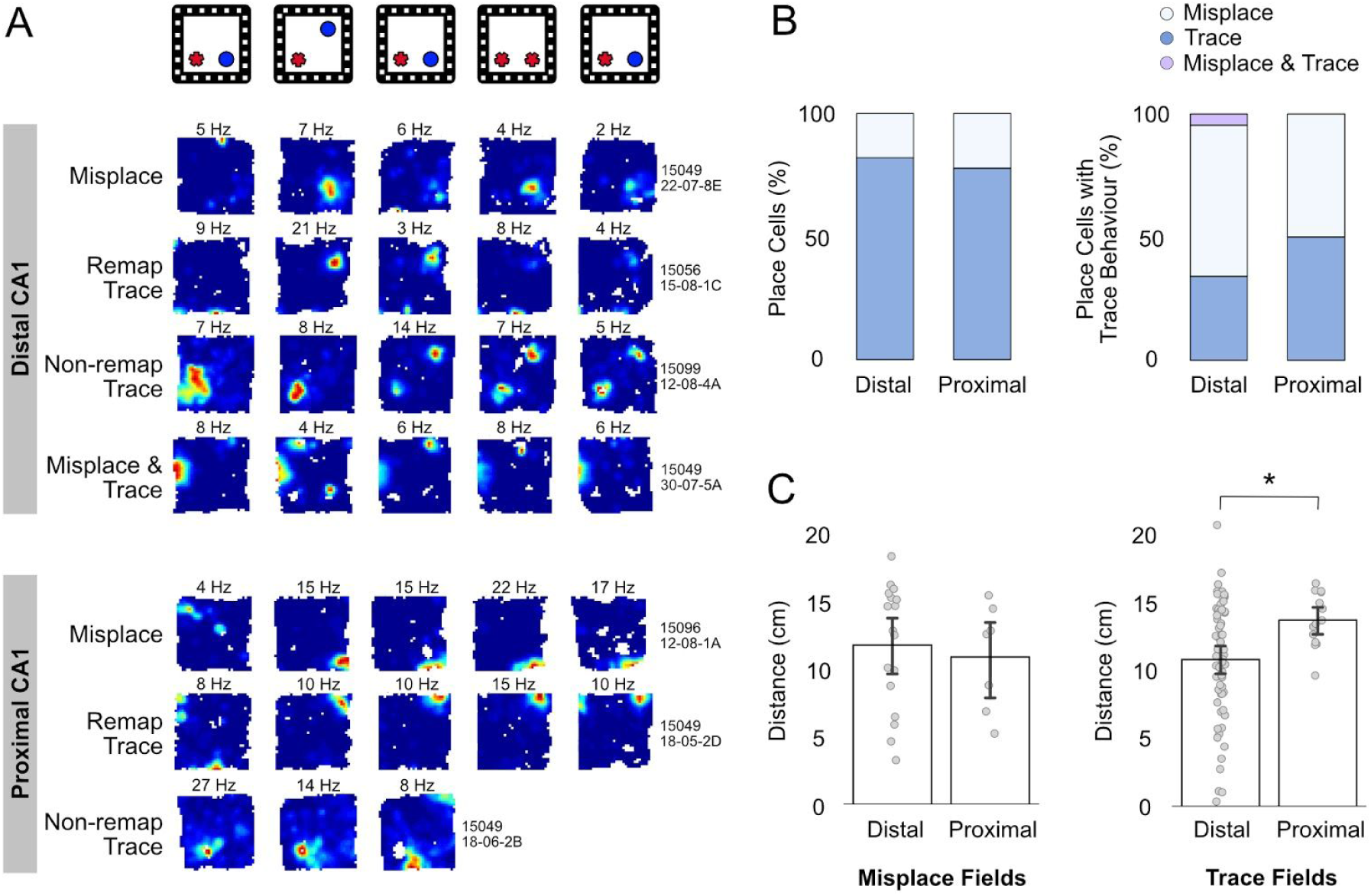
Trace firing across the proximodistal axis of CA1. A) Examples of trace firing in distal (top) and proximal CA1 (bottom). An example of misplace, remap trace, and non-remap trace cells are included for each region. Warm colours indicate high firing rates, and cool colours indicate low firing rates or no firing. Peak firing rates for each trial are indicated above the rate maps. B) Stacked bar charts indicate the proportion of place cells in each region of CA1 with trace firing (left) and the relative proportions of these which conformed to patterns consistent with misplace firing and trace firing at the empty novel object location (right). C) Bar plots show the average distance of place field centroids from the empty object location for misplace (left) and trace cells which fire at the empty novel object location (right). Each grey dot represents a single place field. Asterisk indicates a *p*-value < 0.05 (*).

Given our observation that distal CA1 place cells are more selective for object locations and that remapping in this region is modulated by the proximity of place fields to objects, we hypothesised that trace cells in distal CA1 may more precisely encode object location than trace cells in proximal CA1. To explore this possibility, we measured the distances of place fields expressed in the empty quadrant from the previous object location. For misplace cells, although place fields in distal CA1 were closer to the empty object location than place fields in proximal CA1, this difference was not significant (Figure 5C; *F*_(1, 24)_ = 0.954 *p* = 0.338). However, the place fields expressed by object-place memory cells in distal CA1 were significantly closer to the empty object location than those expressed in proximal CA1 (*F*_(1, 73)_ = 5.586, *p* = 0.021, η^2^ = 0.071). These data demonstrate that LEC inputs drive precision in CA1 place cell representations of previous object locations.

## Discussion

Our data provide evidence that the spatial framework generated by place cells in CA1 is modulated by differential projections from EC in cue-rich environments. Place cells in distal CA1, which receive inputs from LEC, generated more precise representations of objects and locations where objects were previously experienced than place cells in proximal CA1 that are downstream from MEC. Place cells in distal CA1 demonstrated higher spatial tuning and were more stable across trials, particularly when an object was moved to a new location. However, place cell stability in distal CA1 was modulated by the proximity of place fields to objects, where place cells that fired near the displaced object were significantly less stable than place cells with fields elsewhere in the environment. Distance-based modulation of place cell stability was not observed in proximal CA1. Trace cells were recorded in both regions of CA1, but representations of previous object locations were more precise in distal CA1. Overall, our findings suggest that the presence of objects in the local environment drives precision in distal CA1 place cell representations of current object locations and the locations where objects were previously encountered.

These results inform our understanding of information processing within the EC-hippocampal network. Our data suggest that models describing the combination of spatial information from MEC with non-spatial information from LEC within the hippocampus are overly simplistic (Ainge, Tamosiunaite, et al., 2007; Ainge et al., 2012; Ainge, van der Meer, et al., 2007; Eichenbaum et al., 2012; Ferbinteanu & Shapiro, 2003; Hargreaves, Rao, Lee, & Knierim, 2005; Hasselmo, 2009; Hayman & Jeffery, 2008; Kerr, Agster, Furtak, & Burwell, 2007; Knierim, Lee, & Hargreaves, 2006; Leutgeb et al., 2005). Our findings show that the binding of item information into spatial context is not uniform across the hippocampus. These data are consistent with reports of differential functional properties across the proximodistal axis of CA1 (Beer et al., 2018; Hartzell et al., 2013; Henriksen et al., 2010; Ito & Schuman, 2012; Nakamura et al., 2013; Nakazawa et al., 2016) and suggest that the precise integration of item and spatial information necessary for episodic memory may be a specialized function of specific networks within the hippocampus.

Our data also have implications for the use of object information within the EC-hippocampal network. Information about objects is suggested to originate in perirhinal cortex (PRh) (Brown, Warburton, & Aggleton, 2010; Brown & Aggleton, 2001), a structure that is required for novel object recognition (Ennaceur, Neave, & Aggleton, 1996; Mumby & Glenn, 2000; Norman & Eacott, 2005; Wan, Aggleton, & Brown, 1999; Winters, Forwood, Cowell, Saksida, & Bussey, 2004) and contains single neurons that respond to objects (Ahn & Lee, 2015; Bogacz & Brown, 2003; Bogacz, Brown, & Giraud-Carrier, 2001; Burke et al., 2012; Deshmukh, Johnson, & Knierim, 2012). PRh provides major input into LEC but also significant input into MEC and some direct input into hippocampus (Burwell & Amaral, 1998a, 1998b; Furtak, Wei, Agster, & Burwell, 2007; Kosel, Van Hoesen, & Rosene, 1983). It is therefore unlikely that object information reaches the hippocampus from PRh exclusively through connectivity with LEC. Consistent with this, both LEC and MEC encode information about objects in the environment yet manifest distinct patterns of object-modulated firing. LEC neurons generate representations of current object positions and encode locations in the environment where objects were previously experienced (Deshmukh & Knierim, 2013; Tsao et al., 2013, 2018). In contrast, spatial frameworks generated in MEC are tied to environmental stimuli (Chen, Manson, Cacucci, & Wills, 2016; Hafting, Fyhn, Molden, Moser, & Moser, 2005; Pérez-Escobar, Kornienko, Latuske, Kohler, & Allen, 2016) and single cells encode vector relationships between objects and an animal’s position (Høydal et al., 2019). Signalling from object-responsive cells in MEC and LEC might drive different patterns of object modulation across the proximodistal axis of CA1. Our data suggests a model where object information from LEC supports the generation of a spatial map in distal CA1 with higher levels of spatial tuning and precision for object locations, particularly when the positions of objects change. In parallel, object vector signalling from MEC generates place cell representations in proximal CA1 that are less spatially tuned in the presence of objects and less stable across trials.

This suggestion is consistent with previous studies proposing that LEC and MEC support local and global spatial frameworks, respectively (Knierim & Hamilton, 2011; Knierim, Neunuebel, & Deshmukh, 2014; Neunuebel, Yoganarasimha, Rao, & Knierim, 2013). Neunuebel et al. (2013) showed that when local and global cues were put into conflict, the activity of LEC neurons was weakly modulated by local cues, whereas the activity of MEC neurons was modulated by global cues. Consistent with this, Kuruvilla and Ainge (2017) showed that lesions of LEC, but not MEC, impaired rats ability to use local spatial frameworks to guide behaviour. However, anatomical studies suggest a different interpretation of these findings. LEC receives extensive inputs from olfactory areas, whereas MEC receives inputs that carry predominantly visual information (Canto, Wouterlood, & Witter, 2008; Kerr et al., 2007; van Strien, Cappaert, & Witter, 2009). This suggests that representations of environmental stimuli within LEC and MEC are based on different combinations of sensory modalities. In our experiments, objects served as local cues from which rats could sample both visual and olfactory features. Global cues were large objects within the lab from which rats could only extract visual features. Our findings of more precise spatial tuning in place cells that receive LEC input could be driven by those cells having more sensory information (olfactory and visual) from which to construct spatial representations. Whether the distinction between information processing in LEC-MEC is best described in terms of spatial scale (local-global) or sensory modality (olfactory vs. visual) remains to be determined.

Recent studies conceptualise differences in LEC and MEC information processing in terms of egocentric and allocentric spatial frameworks. Wang et al. (2018) analysed spatial frameworks of neurons recorded from LEC and MEC and showed that LEC neurons robustly encoded egocentric space while MEC was more responsive to allocentric cues. Kuruvilla et al. (in press) tested how egocentric and allocentric spatial frameworks support object-location memory in rats. Rats were presented with an object-location memory test from a familiar or novel perspective. Presenting the test environment from a novel perspective forces the rat to orient itself in allocentric space before performing the task, whereas presentation from a familiar perspective encourages the use of an egocentric strategy. Rats with LEC lesions were impaired on the egocentric version, but performed above chance in the allocentric version. Given that egocentric space is governed largely by local cues, this is consistent with the our finding that an LEC-distal CA1 network precisely encodes the location of objects within the immediate environment whereas MEC-proximal CA1 networks encode objects in reference to allocentric spatial frameworks.

Our observations raise further questions regarding how object-related responses in CA1 relate to the memory and navigation functions performed within the EC-hippocampal network. LEC is required to integrate different features of an episode (Chao, Huston, Li, Wang, & de Souza Silva, 2016; Kuruvilla & Ainge, 2017; Rodo et al., 2017; Van Cauter et al., 2013; Vandrey et al., 2020; Wilson, Langston, et al., 2013; Wilson, Watanabe, Milner, & Ainge, 2013), and our data further suggests that the multi-modal representations generated in LEC support place field representations in distal CA1 that precisely encode items within an environment and are sensitive to changes in their position. Further, the precise representations of previous object locations in distal CA1 suggest a role of LEC-hippocampus circuitry in object-place memory. Our findings are consistent with a role of projections from LEC to the hippocampus in episodic memory (Vandrey et al., 2020) and with reports that LEC manifests early pathology in Alzheimer’s disease (Gómez-Isla et al., 1996; Khan et al., 2014; Kobro-Flatmoen, Nagelhus, & Witter, 2016; Stranahan & Mattson, 2010). In contrast, MEC is part of a network that supports path integration, navigation, and spatial memory (Hales et al., 2014; Steffenach, Witter, Moser, & Moser, 2005; Tennant et al., 2018; Van Cauter et al., 2013). Our data suggest that spatial representations generated in MEC drive spatial representations in proximal CA1 that are less tuned to local features. This is consistent with the utility of non-local cues for navigation and spatial memory, where global cues provide consistent spatial information when navigating over longer distances.

## Acknowledgements

This work was supported by a Henry Dryerre scholarship from the Royal Society of Edinburgh to B.V.

## Author Contributions

Conceptualization and Methodology, B.V., J.A., Investigation, B.V., Writing -Original Draft, B.V., J.A., Writing, Review & Editing, B.V., J.A., Supervision, J.A., Funding Acquisition, B.V.

---

**Figure S1:**
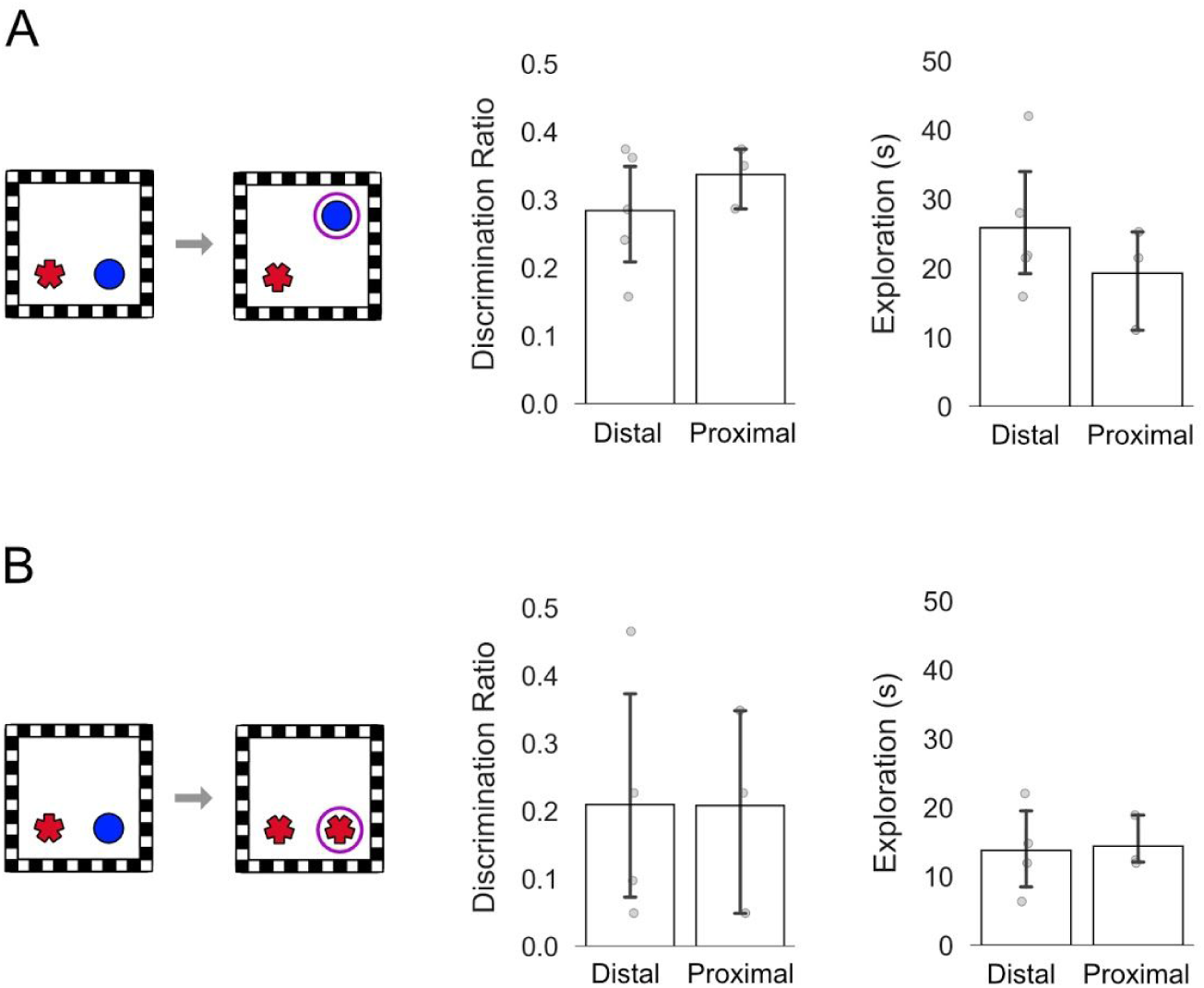
Animals in both groups can discriminate novel object positions and identities. (Left) Schematics show the spatial change that occurs in the object displacement trial (A) and object-place recognition trial (B). Pink circle indicates the object that is in a novel configuration. Bar graphs show average discrimination ratios (left) and time spent exploring the objects (right). Discrimination ratios are calculated as the amount of time spent exploring the object in a novel configuration subtracted by the amount of time spent exploring object in a familiar configuration, divided by the total exploration time [36]. A positive discrimination ratio indicates a preference for the object in the novel configuration. Animals in with implants targeting distal and proximal CA1 explored the novel configuration more than predicted by chance in the object displacement (distal CA1: *t*(4) = 7.487, *p* = 0.002, proximal CA1: *t*(2) = 9.831, *p* = 0.005) and object-place recognition trial (distal CA1: *t*(4) = 2.978, *p* = 0.041, proximal CA1: *t*(2) = 3.306, *p* = 0.040). There was no significant difference in either task in recognition of the novel configuration (stats) or total exploration (stats). Each grey dot represents a value for an individual animal. Note, one animal was implanted with electrodes targeting distal and proximal CA1. Error bars are SEM.

**Figure S2:**
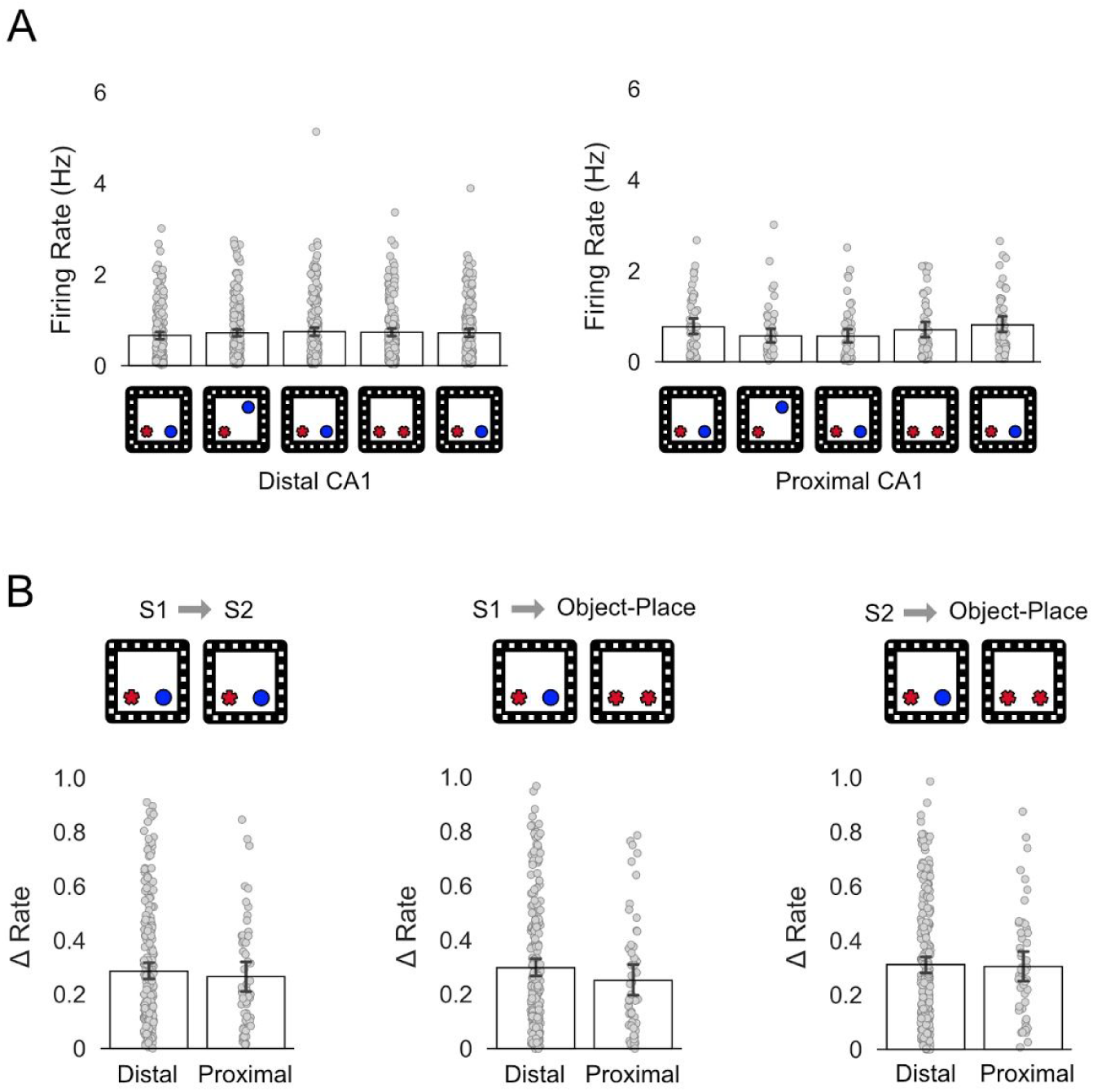
Firing rates of distal and proximal CA1 place cells in response to changes in object identity. A) Bar chart shows average firing rates of place cells in distal CA1 (left) and proximal CA1 (right) across all trials. B) Changes in firing rate (Δ) between the first two standard trials (left), the first standard trial (S1) and the object-place recognition trial (middle), and the second standard trial (S2) and the object-place recognition trial (right). Δ was calculated by finding the absolute value of the firing rate difference between the two trials, and dividing this value by the sum of firing rates across the two trials (Lu et al., 2013). There was no significant difference between firing rate changes observed in distal and proximal CA1 across S1 and S2 (F (_1, 290_) = 0.3261, *p* = 0.568), S1 and the object-place recognition trial (*F* (_1, 274_) = 0.0378, *p* = 0.846) and S2 and the object place recognition trial (*F* (_1, 274_) = 1.0715, *p* = 0.3015). Grey dots represent values for single place cells. Error bars are SEM.

